# Hidden drivers of restoration: Persistent divergence in soil microbiome functional capacity post-habitat reconstruction

**DOI:** 10.64898/2026.08.31.747609

**Authors:** Vicki W. Li, Alma L. Reyes, Kasey N. Kiesewetter, Eric Cline, Amanda H. Rawstern, Carlos Coronado-Molina, Fred H. Sklar, Michelle E. Afkhami

**Author notes:** Correspondence: Vicki W. Li, Department of Biology, University of Miami, Coral Gables, FL 33146, USA.

## Abstract

Ecosystems today are facing unprecedented environmental stress, leading to large-scale losses of habitat and ecosystem services. To address this, reconstructive efforts aim to restore habitat features and their natural complexity, biodiversity, and function. However, many reconstructive efforts fail to consider microbial communities, even though they play key roles in decomposition, nutrient cycling, and plant and animal health. Here, we use shotgun metagenomic sequencing to compare the structure and functional capacity of soil microbial communities from natural Everglades tree islands and islands constructed within a landscape-scale experimental Everglades restoration effort, followed by a manipulative greenhouse experiment to link tree sapling traits with microbiome functional genetic divergence. We found that constructed and natural island microbiomes exhibited strong, robust, and persistent divergence in both taxonomic and functional composition, with natural island microbial communities having greater functional genetic diversity and redundancy. Furthermore, we identified enrichment in functional pathways in constructed island microbiomes such as those involved in pollutant degradation that may reflect continued disturbance leading to shifts in microbiome functional profiles. Despite their capacity for functions such as nitrogen cycling we found to be important for supporting sapling growth, constructed island microbiomes displayed reduced functional genetic diversity and redundancy and were enriched in pathways associated with environmental disturbance, suggesting a potentially diminished capacity for long-term resilience. Overall, our assessment of soil microbial communities in reconstructed and natural habitats emphasizes how reconstructive restoration can impact microbial functional repertoires and highlights the importance of these hidden players in management and restoration of ecosystem health.

## Introduction

Human activities have led to widespread habitat loss and degradation, contributing to tremendous declines in biodiversity and ecosystem services (Díaz et al., 2018; Eyres et al., 2025). To address this, ecological restoration aims to improve habitat quality and ultimately recover biodiversity and ecosystem function (Wortley et al., 2013). However, current restoration efforts, particularly those involving reconstructing landscape features, largely focus on macroscopic ecosystem components and therefore fail to consider the impact of restoration practices on microbial communities as well as the role of these microbial communities in the restoration process and/or their restored habitats (Contos et al., 2025; Kiesewetter et al., 2025). Microbes are ubiquitous in ecosystems and perform crucial functions such as decomposition, nutrient cycling, and pollutant degradation (Hartmann & Six, 2023; Wang et al., 2024). Therefore, microbes are relevant, if not central, to many restoration goals and outcomes, both for supporting targets—such as improved ecosystem function and interactions with other organisms (e.g., primary producers)—and also as a restoration goal themselves (Junker, 2026; Singh Rawat et al., 2023).

Although restoring microbial community function and interactions with macroscopic communities is central to broader ecosystem restoration (Contos et al., 2025; Kumaresan et al., 2017), how different restoration approaches impact microbial communities is unclear. For instance, when entire landscape features are reconstructed, initial microbial communities in building materials and substrate may be poorly adapted to their new environments or lack the necessary functional repertoires to underpin macroscopic biodiversity and ecosystem function (Kiesewetter et al., 2025). Alternatively, novel microbiomes in restored habitats may cross thresholds into fully functional “alternative” states, making recovery towards relatively undisturbed community states found in natural habitats unfeasible or even undesirable (Hart et al., 2020). Although the role of microbiomes in restoration is understudied, microbiome management is a promising avenue for restoration because it can have broader or long-term consequences for microbiome function and species interactions (Contos et al., 2025; Kumaresan et al., 2017). Therefore, greater investigation of microbiomes and their functional profiles in reconstructed habitat features is necessary for integrating microbial communities into restoration practices.

In this study, we address this gap by evaluating how large-scale habitat reconstruction impacts microbiome functional genetic repertoires and how associated functional gene shifts impact native tree sapling performance under different hydrological regimes in the iconic Everglades ecosystem, the focus of the most expansive and costly restoration effort in North America (Clarke & Dalrymple, 2003). As the largest subtropical wetland in the United States, the Everglades is host to tremendous biodiversity and helps provide humans with both flood control and water for consumptive use (Brown et al., 2006; Perry, 2008; Richardson, 2010). However, the Everglades has declined to only half of its original extent as a result of human development, habitat destruction, and hydrological management (Perry, 2008; Sklar et al., 2001). In particular, tree islands, aggregations of woody vegetation on elevated peat or limestone, have declined precipitously, by 67% across the entirety of Water Conservation Area 3 (one of three sections of the remaining Everglades outside of Everglades National Park and Big Cypress National Preserve) and by up to 87% in some areas (Sklar & van der Valk, 2002; Wetzel et al., 2005). These tree islands are nutrient and biogeochemical hotspots in an otherwise oligotrophic landscape and are estimated to hold up to two-thirds of total phosphorus in the Everglades system, with soil phosphorus levels on tree islands up to 100 times higher than in surrounding marshes and sloughs (Wetzel et al., 2005, 2009). Therefore, tree island decline, influenced by human modifications of hydrology, leads to loss of habitat, diminished landscape complexity, and release of nutrients into the surrounding landscape, with cascading impacts on surrounding systems adapted to low nutrient conditions (Richardson, 2010; Wetzel et al., 2005). As a result, tree islands are a key priority for Everglades restoration, with efforts to construct new islands from degraded limestone levees currently ongoing.

This reconstructive restoration can have pervasive impacts on microbial communities. Recent work identified taxonomic differences in soil microbiomes between constructed and natural tree islands. Specifically, taxonomic community composition of soil fungi differed between reconstructed and natural Everglades tree islands even 18 years post-island construction (Kiesewetter et al., 2025). Moreover, soil microbiomes from the constructed islands had differential impacts on native tree species, with enhanced benefits under wetter experimental conditions (Kiesewetter et al., 2025). This demonstrates that microbes in reconstructed habitats may persistently differ from their natural counterparts despite largely similar environmental conditions, especially in the absence of active microbial management. Furthermore, there may be functional divergence between taxonomically distinct microbiomes that are responsible for these differential higher-order effects such as interactions with tree communities, but what shifts in microbiome functional repertoires may underlie altered community functions remain largely unknown.

Here, we combine field surveys, shotgun metagenomic sequencing of soil microbial communities, and a manipulative greenhouse experiment to understand functional differences between soil microbiomes from constructed and natural tree islands. Specifically, we evaluate (1) how soil microbial communities differ between constructed and natural tree islands in terms of their taxonomic and functional genetic composition and diversity, functional genetic redundancy, and functional pathways and (2) how these differences in microbiome functional repertoires between constructed and natural tree islands in turn impact native tree sapling performance under different hydrological management regimes proposed for this ecosystem. Answering these questions provides insight into the long-term functional consequences of reconstructive restoration and hydrological management for soil microbiomes.

## Methods

### Study system

The Everglades is an iconic wetland ecosystem consisting of a mosaic of habitats, including tree islands, which cover <5% of its extent but are critical nutrient hotspots in an otherwise naturally oligotrophic landscape (Richardson, 2010; Wetzel et al., 2005). When tree islands decline or are lost, nutrients are released into the surrounding habitat (Richardson, 2010; Wetzel et al., 2005). Tree islands are often teardrop-shaped, with the head (front) of the island having the highest elevation and the downstream tail (back) having lower elevation (Sklar & van der Valk, 2002). The wet hydroperiod peaks in October/November with high water levels (i.e., water stage) that typically inundate island tails and sometimes even entire islands. The dry hydroperiod usually reaches the lowest water levels in May, and as water levels gradually recede, soil surfaces of many tree islands eventually dry out (Sklar & van der Valk, 2002).

To understand the impacts of tree island construction, restoration, and hydrological management, the Loxahatchee Impoundment Landscape Assessment (LILA) was established in 2003 by the South Florida Water Management District and the Army Corps of Engineers in the US Fish & Wildlife Service’s Arthur R. Marshall Loxahatchee National Wildlife Refuge in Palm Beach County, Florida, USA (26.489° N, 80.219° W). Spanning 80 acres, this experimental Everglades landscape consists of eight ∼2500 m^2^ constructed tree islands surrounded by slough and sawgrass ridges, allowing researchers to study how restoration and management impact biodiversity, including microbial communities (Aich et al., 2011; Almeida et al., 2023; Kiesewetter et al., 2025; Reyes et al., *in prep*). Tree islands were constructed with peat or limestone cores, covered with a layer of peat substrate, and planted with a mixture of ten tree species found on tree islands (Stoffella et al., 2010). These constructed tree islands enable us to evaluate how reconstructive restoration approaches without active microbial management shape microbiomes.

### Microbiome sample collection

In October 2021, we collected soil samples from the eight LILA constructed tree islands and 14 nearby natural tree islands located in the Water Conservation Area 3A selected to most closely match the hydroperiods of the LILA tree islands. For each island, we aseptically collected 50 mL soil from both the head and the tail (with the exception of one natural tree island, for which the tail was inaccessible). From each LILA tree island and seven of the natural tree islands, we collected an additional 5 L of soil to be used as inoculum in our greenhouse experiment. Soil was transported back to the University of Miami (Coral Gables, FL) on ice on the same day as collection. Soils for microbial community sequencing were stored in a –20°C manual defrost freezer, whereas inoculum soil was briefly stored at 4°C during establishment of the greenhouse experiment.

### Microbiome DNA extractions, library preparation, sequencing, and bioinformatics

Microbial DNA was extracted from 0.5 g of soil from each sample (N = 42) using a DNeasy PowerSoil Pro Kit (Qiagen, 47016) following standard manufacturer’s protocols for processing soil samples with high water content. Samples were then purified using an E.Z.N.A. Gel Extraction Kit (D2500-02; Omega Bio-tek, Norcross, GA, USA). DNA concentrations were checked with a Qubit 4 Fluorometer (Invitrogen, Q33327). We prepared metagenomic libraries using the Illumina DNA Prep Kit (20060059) with Illumina DNA/RNA Tagmentation Indexes. Metagenomic libraries were sent to the University of Miami’s John P. Hussman Institute for Human Genomics Sequencing Core Facility (RRID:SCR_017828) on the Illumina NovaSeq X Plus (150 bp paired end).

The resulting demultiplexed sequence data was processed using *fastp* (S. Chen et al., 2018) to remove Nextera adapters and low-quality sequences using default settings (phred quality ≥ Q15, unqualified bases limit = 40%); reads shorter than 50 bp were also removed (see Broderick et al., 2025). We checked processed reads for quality using *FastQC* (Andrews, 2023) and performed error correction using *bbcms* (*BBTools*; Bushnell, 2014). Reads were assembled into contigs within each metagenomic sample using *metaSPAdes* (Bankevich et al., 2012; Nurk et al., 2017), retaining only contigs ≥500 bp using *reformat* (*BBTools*; Bushnell, 2014). Subsequent functional annotation of contigs for KEGG KOs (Kanehisa et al., 2016) occurred using the MOSHPIT toolkit on the QIIME2 platform (Bolyen et al., 2019; Ziemski et al., 2025), with *eggnog-mapper v2* (Cantalapiedra et al., 2021; Huerta-Cepas et al., 2019; Hyatt et al., 2010) alongside the *DIAMOND* aligner (Buchfink et al., 2021). We extracted functional annotations with an e-value cutoff of 0.0001. For taxonomic characterization of microbial communities, filtered reads were classified using Kaiju v1.8.0 using default parameters and NCBI’s *nr_euk* database (Menzel et al., 2016).

### Plant–microbiome experiment

To understand how microbiomes from constructed and natural tree islands in conjunction with hydrological management affect native sapling performance, we conducted a factorial greenhouse experiment manipulating microbiome origin (constructed versus natural tree island source), inoculation (live versus sterilized soil inoculum), and hydrological treatment (unconstrained versus constrained) across four tree species commonly found on tree islands and planted in restoration. The tree species included in our experiment were *Eugenia axillaris* (Swartz) Willdenow (white stopper), *Ilex cassine* L. (dahoon holly), *Annona glabra* L. (pond apple), and *Chrysobalanus icaco* L. (cocoplum). Saplings were sourced from a native nursery (Indian Trails Native Nursery in Lake Worth, FL), where they were propagated from seeds produced by local Everglades genetic stock.

For each sapling, we nested the pot in larger vessels to allow manipulation of hydrological treatment (design modified from Almeida et al., 2023). We added inoculum (3% of pot volume) of live or sterile soil collected from one of 15 islands. Live soil inoculum contained a microbial community from one of the sequenced tree islands (see Appendix: Section S1 *Microbiome DNA extractions, library preparation, and sequencing* and Appendix: Table S1 for details), whereas sterile inoculum consisted of the same soils autoclaved three times at 121°C. “Unconstrained”, or wet, and “constrained”, or dry, hydrological treatments were applied to test microbial effects on plants under different hydrological regimes (see Appendix: Section S1 *Manipulation of hydrological treatments in the plant–microbiome experiment* for methodological details).

Saplings were grown in the University of Miami Greenhouse (Coral Gables, FL). All saplings were allowed to first acclimate to greenhouse conditions. Afterwards, microcosms were established and initially watered daily to fill them to appropriate water levels, then watered every two days to maintain their water levels. We recorded trunk diameter and leaf number at five months (roughly equivalent to the length of the wet season, a major hurdle for success of new plantings). We measured stomatal conductance with the LI-600 Porometer (LI-COR Biosciences, Lincoln, NE) on all saplings with at least one healthy leaf at the end of the experiment (two weeks after sapling performance data collection at five months).

### Data analysis

All statistical analysis was performed in R version 4.5.2 (R Core Team, 2025). To characterize taxonomic differences between microbiomes from constructed and natural tree islands, we used our Kaiju taxonomic classifications to the genus level to perform a permutational multivariate analysis of variance with 999 permutations (PERMANOVA in *vegan*; Oksanen et al., 2026) with island type (constructed versus natural) and location (head versus tail) as explanatory terms, enabling us to determine whether there were significant differences in taxonomic composition. We did so using both robust Aitchison (based on relative abundances of microbial genera) and Jaccard (based on genera presence versus absence) distance matrices. We also performed linear mixed-effects models using *lme* (*nlme*; Pinheiro & Bates, 2025) to test whether taxonomic diversity and richness differed by island type, with taxonomic Shannon-Wiener diversity or richness as the response variable and the same explanatory variables as in our PERMANOVA as well as island as a random effect. To characterize differences in functional repertoires between microbiomes from constructed and natural tree islands, we repeated these analyses for functional gene (i.e., KEGG KO) composition, diversity, and richness.

Beyond differences in overall functional repertoires, communities can differ in their functional redundancy. Functional redundancy measures the degree to which taxa in a community overlap in their functional abilities, and is thought to enhance microbiome stability by allowing for losses of taxa without broader losses of community function (Afkhami et al., 2026). To evaluate differences in functional redundancy between constructed and natural tree islands, functional redundancy for each KEGG functional pathway within each sample was calculated using Kaiju taxonomic assignments to the genus level as

FR_ij_ = –Σ*Pi*ln*Pi*

where *P_i_* represents the relative frequency of functional pathway *i* encoded by a specific genera in a sample *j* compared to the total number of encoded functional pathways in the sample (Li et al., 2024). Furthermore, because some functional pathways may generally exhibit greater degrees of functional redundancy than others regardless of source microbial community, we additionally employed a paired design when testing for differences in functional redundancy between constructed and natural islands. Specifically, differences in functional redundancy were evaluated using a paired t-test comparing the difference in mean functional redundancy across constructed islands compared to the mean functional redundancy across natural islands for each given KEGG functional pathway. Functional pathways that were absent from all constructed island samples or all natural island samples could not be compared and were therefore excluded, representing only 0.7% of functional pathways included in this analysis (3 pathways out of 437 total). We note that these functional redundancy results were robust to different statistical approaches (for details see Appendix: Section S1 *Constructed and natural tree island microbiomes show robust differences in functional genetic redundancy*).

To further investigate which specific functional genes were more abundant in constructed or natural islands, we used ANCOM-BC2 with the default prevalence cutoff of 0.10 (H. Lin & Peddada, 2024) to identify genes that exhibited differential abundance in either constructed or natural tree island microbiomes. Based on our ANCOM-BC2 output, we performed gene set enrichment analysis (GSEA) with Benjamini-Hochberg correction using the *gseKEGG* function (*clusterProfiler*; Yu et al., 2012) to determine whether certain functional pathways were enriched in soil metagenomes in one island type (constructed/natural) compared to another.

To explore the impact of microbiome functional potential and hydrology on native saplings, we evaluated how island type, hydrological treatment, and microbiome presence affected sapling trunk diameter (woody growth), leaf number (foliar growth), and stomatal conductance (physiological response). We ran linear mixed-effects models using *lmer* (*lme4*; Bates et al., 2026) for each response metric with island as a random effect and the following fixed effect explanatory variables: inoculation treatment (sterile versus live), water treatment (constrained versus unconstrained), tree species identity, island type (constructed versus natural), the primary axis of variation in microbiome functional gene (KEGG KO) composition (using Aitchison distance matrices) from the multivariate principal coordinate analysis (PCoA) described above, functional gene diversity, and functional pathway redundancy. In addition, we included all possible interactive effects between inoculation treatment, hydrological treatment, and tree species identity, as well as the interactions between each of those terms and with island type, the primary axis of microbiome functional gene composition, functional gene diversity, and functional pathway redundancy. We chose to use the primary axis of variation in functional gene composition to describe microbial communities because it explained the greatest amount of variation in composition and because constructed and natural tree island communities diverged significantly along this axis (χ^2^_1_= 5.3087, df = 1, p = 0.02). Furthermore, we used the axis based on Aitchison rather than Jaccard distance matrices because the resulting primary axis of variation explained a greater proportion of total variance (Aitchison: 53.6% versus Jaccard: 14.2%). Following the construction of our global models for each sapling response metric, we performed global model selection based on the corrected Akaike Information Criterion (AIC_c_) using the *dredge* function in R (*MuMIn*; Bartoń, 2024). Because sapling species identity and interactions with species were not retained in the best model for any of our sapling response metrics, we did not run models for each species individually. Additionally, because our best model for trunk diameter retained the primary axis of variation in functional gene composition as an explanatory term, we further inspected this axis by using the *envfit* function in R (*vegan*; Oksanen et al., 2026) to correlate KEGG KO relative abundances to PCoA axes. We then used a stringent approach to filtering correlated genes to avoid retaining extraneous genes. Specifically, after retaining only KOs highly significantly correlated to the primary axis following Benjamini-Hochberg correction (adjusted p < 0.01), we further filtered our list of functional genes to only retain those that were strongly correlated to the primary axis of variation in functional gene composition (r ≥ 0.7 for the primary axis) as well as well-explained overall by our PCoA (R^2^ ≥ 0.7). We then performed Boruta feature selection with 999 permutations (*Boruta*; Kursa & Rudnicki, 2026) to identify which filtered KEGG KOs were predictive of the primary axis of variation in functional gene variation, producing a final curated list of 30 particularly relevant functional genes (Table S2). We manually inspected the function of these 30 KEGG KOs to gain insight into the primary axis of variation from our PCoA.

## Results

We identified robust differences in both functional gene and taxonomic composition between constructed and natural tree islands, demonstrating persistent and long-term divergence of microbial communities in reconstructed habitats from their natural counterparts. Furthermore, natural tree island microbiomes exhibited higher functional gene diversity and functional pathway redundancy. Both constructed and natural tree island microbial communities were enriched for subsets of functional genes and their associated functional pathways, with natural tree island communities relatively enriched in pathways related to general cellular functions and carbohydrate metabolism, and constructed island communities enriched in pathways related to lipid metabolism, pollutant degradation, and pathogenicity or antibiotic resistance. Microbiome functional gene composition was also linked to sapling growth, with more negative values along the primary axis of variation corresponding to greater trunk diameter. These negative values generally corresponded more to constructed tree island microbiomes and were correlated with multiple genes involved in nitrogen cycling.

### Constructed and natural tree island microbiomes show long-term differences in community composition

Even 18 years post-reconstruction, constructed and natural tree island microbial communities differed significantly in community composition at the genus level, both by relative abundance of genera (Aitchison: F_1,39_ = 13.29, p = 0.001; Figure 1a) and by genera presence versus absence (Jaccard: F_1,39_ = 1.99, p = 0.002; Figure 1b). No differences in taxonomic diversity (χ^2^_1_= 0.44, p = 0.51) or richness (χ^2^_1_= 0.62, p = 0.43) between constructed and natural tree island communities were identified. Additionally, no significant differences in taxonomic composition by either distance metric (Aitchison: F_1,39_ = 2.06, p = 0.12; Jaccard: F_1,39_ = 1.03, p = 0.27), diversity (χ^2^_1_= 1.15, p = 0.28), or richness (χ^2^_1_= 1.83, p = 0.18) were identified based on island location (i.e., head versus tail).

**Figure 1.**
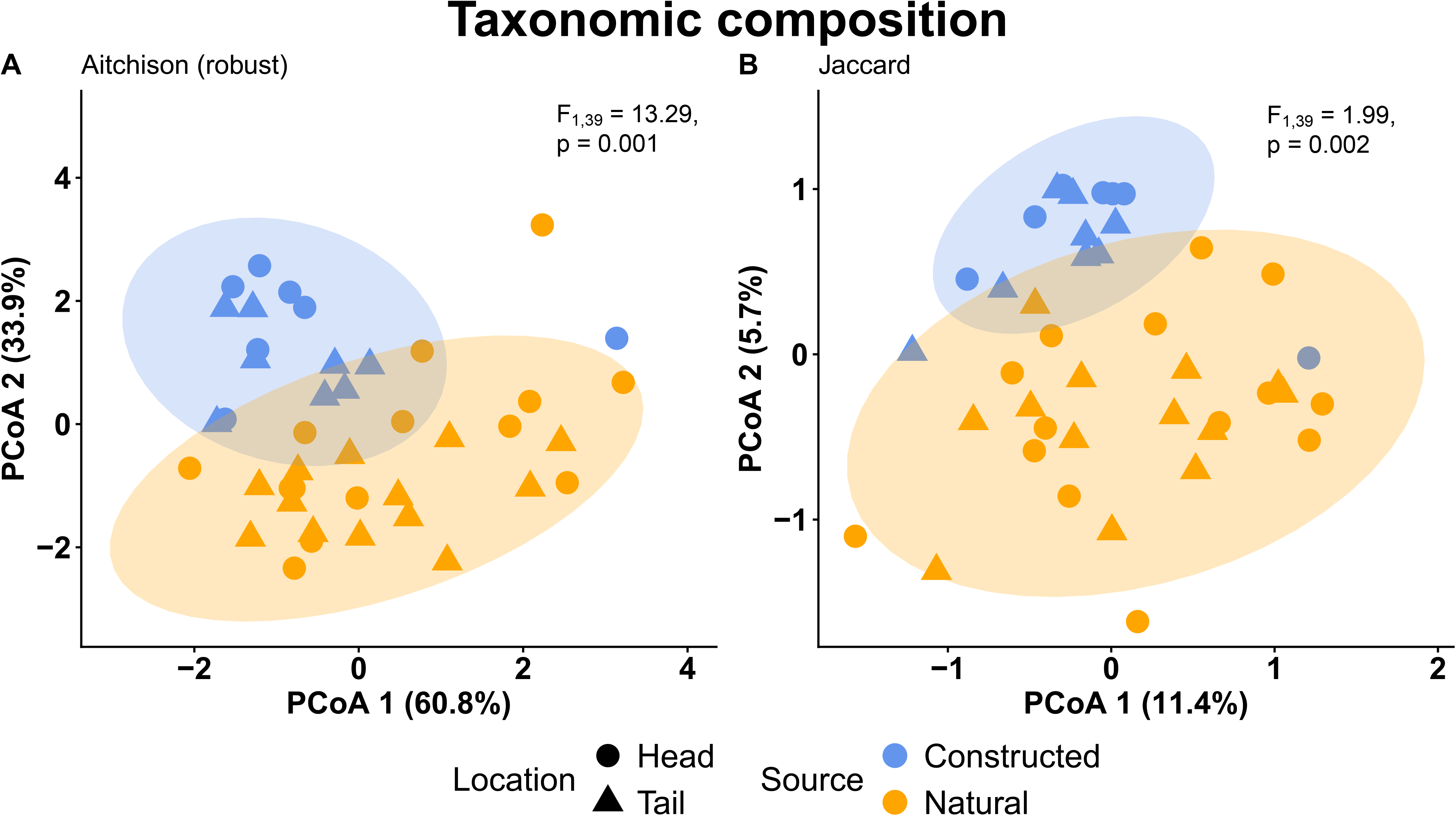
Microbial community taxonomic (at the genus level) composition differed between constructed (blue) and natural (orange) tree islands based on both **(A)** Aitchison (PERMANOVA: F_1,39_ = 13.29, p = 0.001) and **(B)** Jaccard (PERMANOVA: F_1,39_ = 1.99, p = 0.002) distance matrices.

### Identified taxonomic differences are reflected in differences in functional repertoires

Taxonomic differences between constructed and natural tree island microbiomes were reflected in differences in functional repertoires, with functional gene composition differing significantly by island type (Aitchison: F_1,39_ = 14.03, p = 0.001; Jaccard: F_1,39_ = 3.2879, p = 0.001; Figure 2). Furthermore, natural tree island microbiomes exhibited slightly but significantly greater functional gene diversity than constructed island microbiomes (χ^2^_1_= 5.65, p = 0.017; constructed: 7.80 ± 0.01 versus natural: 7.83 ± 0.01; mean ± S.E.M.; Figure 3a), although functional gene richness was similar (χ^2^_1_= 0.01, p = 0.91). Location of sampling on island heads versus tails did not significantly affect functional gene composition (Aitchison: F_1,39_ = 0.49, p = 0.60; Jaccard: F_1,39_ = 0.98, p = 0.41), diversity (χ^2^_1_= 0.62, p = 0.43), or richness (χ^2^_1_= 0.12, p = 0.73), suggesting that available functional gene repertoires are similar at an island-level scale.

**Figure 2.**
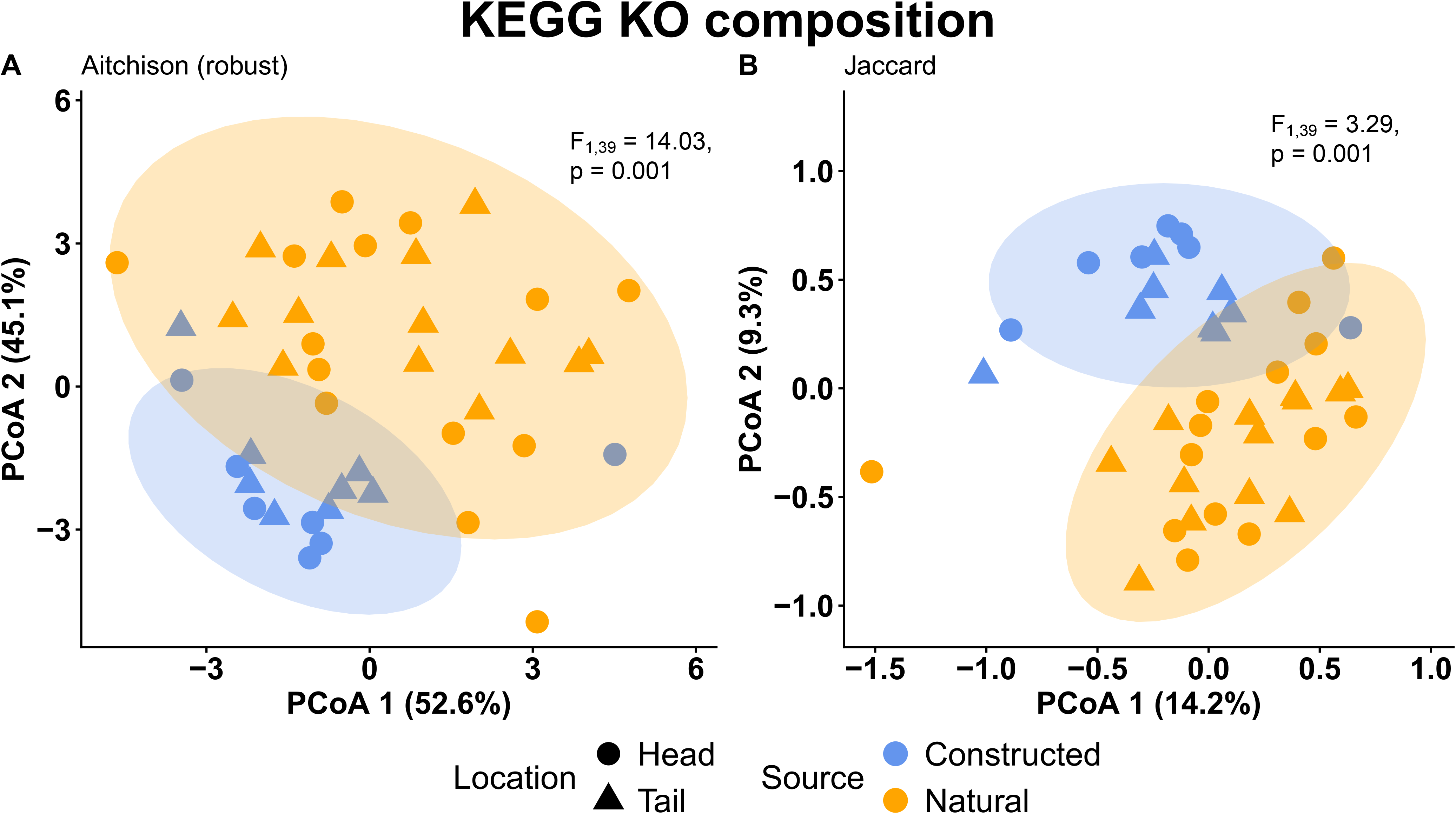
Microbial community functional gene (KEGG KO) composition differed between constructed (blue) and natural (orange) tree islands based on both **(A)** Aitchison (PERMANOVA: F_1,39_ = 14.03, p = 0.001) and **(B)** Jaccard (PERMANOVA: F_1,39_ = 3.29, p = 0.001) distance matrices.

**Figure 3.**
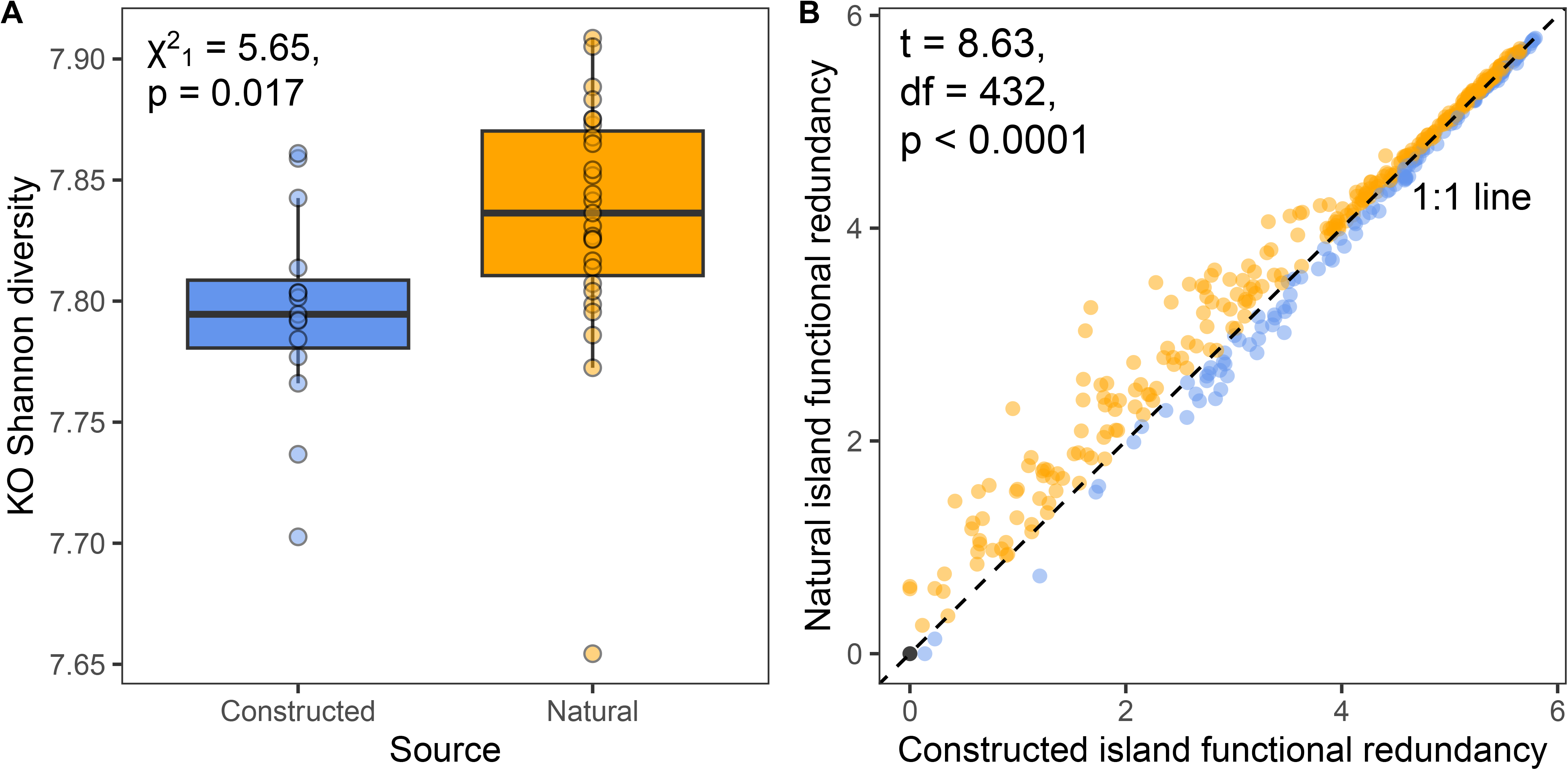
(A) Natural tree island soil microbiomes (orange) exhibited relatively greater functional gene (i.e., KEGG KO) Shannon diversity than constructed island microbiomes (blue; linear mixed-effects model: χ^2^_1_= 5.65, p = 0.017). **(B)** Natural island microbiomes also exhibited greater functional (KEGG) pathway functional redundancy than constructed island microbiomes (paired t-test: t = 8.63, df = 432, p < 0.0001). For visualization, each unique functional pathway is represented by a point, with its average functional redundancy across constructed tree island communities along the x-axis and its average functional redundancy across natural island communities on the y-axis. Pathways (i.e., points) with relatively greater functional redundancy in constructed islands are depicted in blue and occur below the 1:1 line, whereas those with greater functional redundancy in natural islands are shown in orange above the 1:1 line. Those with equivalent functional redundancies between constructed and natural tree island communities are shown in black. Functional pathways that were entirely absent across all constructed or all natural island communities were excluded from this analysis (n = 3).

After identifying differences in functional repertoires between constructed and natural tree island microbiomes, we then investigated whether these functional abilities exhibited differences in community-level redundancy. When evaluating KEGG functional pathways encoded by microbial contigs assigned to the genus level, we found that overall, natural tree island microbial communities had higher functional redundancy (paired t-test: t = 8.63, df = 432, p < 0.0001; mean ± S.E.M. difference in functional redundancy for a given functional pathway: 0.11 ± 0.01; n = 433 pathways; Figure 3b).

To investigate the implications of the differences in functional repertoires we identified, we used ANCOM-BC2 to identify functional genes (i.e., KEGG KOs) that were differentially abundant between constructed and natural tree island microbial communities. In total, 34 functional genes exhibited statistically significant differential abundance, with 11 enriched in constructed island microbiomes and 23 enriched in natural islands (see Appendix: Figure S1). Upon performing GSEA, 37 functional (KEGG) pathways were found to be significantly differentially enriched, with 23 enriched in constructed tree islands and 13 in natural tree islands. Functional pathways enriched in natural tree island microbiomes were often associated with general cellular functions or carbohydrate metabolism, whereas those enriched in constructed island communities were often related to lipid metabolism, pollutant degradation, or pathogenicity and antibiotic resistance (Figure 4).

**Figure 4.**
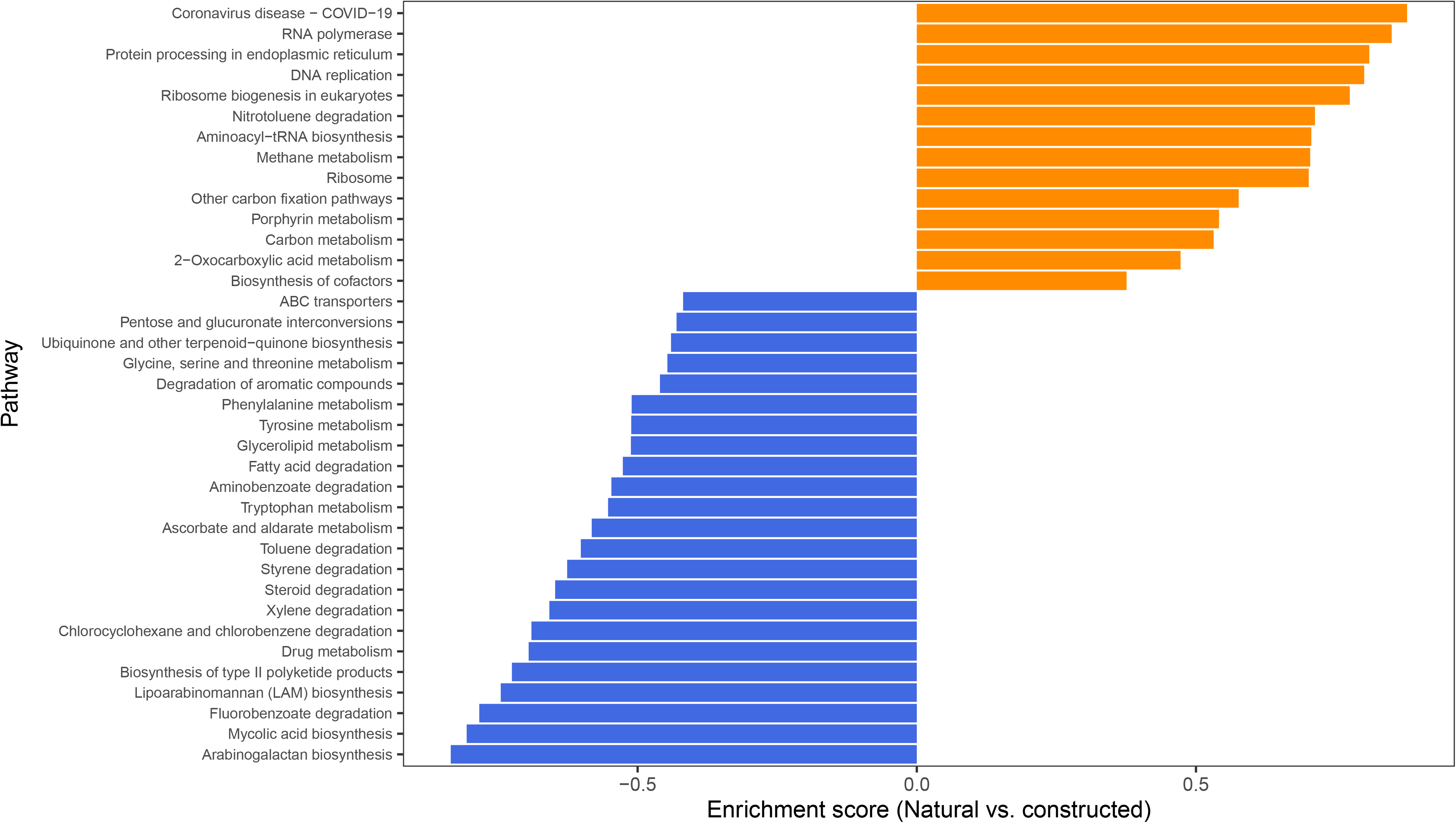
Gene set enrichment analysis of functional (i.e., KEGG) pathways relatively enriched in constructed (blue; negative enrichment score) and natural (orange; positive enrichment score) tree island soil microbial communities.

### Functional genetic repertoires of soil microbiomes explain variation in microbial effects on tree sapling growth

Woody growth of saplings in our experiment was significantly affected by variation in microbiome functional genetic repertoires and hydrological regimes. Specifically, the best supported model for trunk diameter retained hydrological treatment (constrained/unconstrained) and the interaction between microbial treatment (live/sterile inoculum) and the primary axis of variation in microbiome functional gene composition as fixed effect explanatory terms. Importantly, we found trunk diameters were larger in the constrained (dry) treatment (χ^2^_1_= 17.63, df = 1, p < 0.0001; Figure 5a). Furthermore, in the live inoculum treatment, inocula with more negative values along the primary axis of functional gene variation supported larger trunk diameters (χ^2^_1_= 7.90, df = 1, p = 0.0049; Figure 5b). In contrast, leaf number and stomatal conductance did not respond to differences in constructed versus natural islands (i.e., the null model was the best-supported model based on global model selection), suggesting that unlike trunk diameter, functional genetic variation in microbiomes was less important for these sapling response metrics. (See Appendix: Tables S14–S16 for more details.)

**Figure 5.**
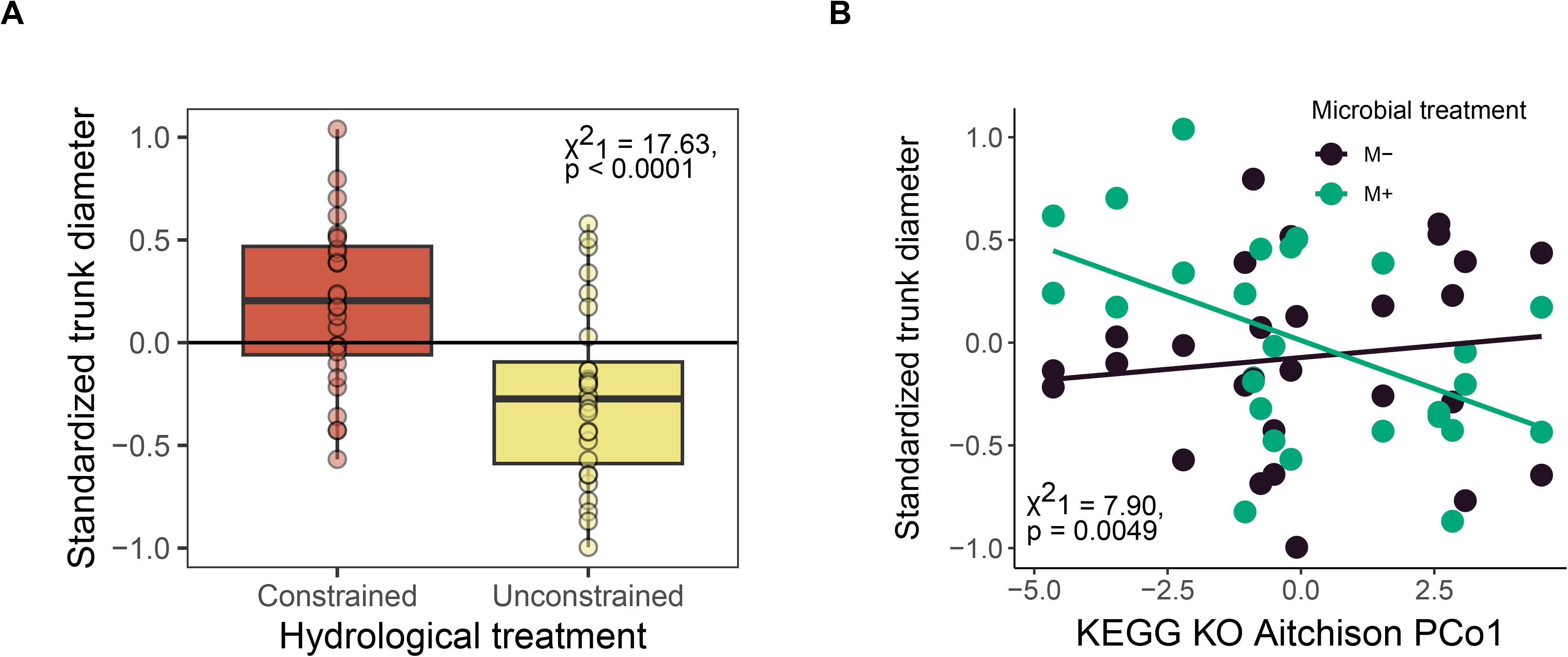
Standardized trunk diameter, a metric for woody growth, was **(A)** generally higher in the constrained (dry) hydrological treatment (χ^2^_1_= 17.63, df = 1, p < 0.0001), and also **(B)** depended on the interaction between microbial treatment (live/M+ versus sterile/M– inoculum) and the primary axis of variation in microbiome functional gene composition based on Aitchison distance matrices (χ^2^_1_= 7.90, df = 1, p = 0.0049). For the live microbial treatment, more negative values along the primary axis of variation contributed to larger trunk diameters.

Constructed and natural tree island communities diverged significantly along the primary axis of functional gene composition (χ^2^_1_= 5.3087, df = 1, p = 0.02), with more negative values along this axis generally corresponding to constructed tree island microbiomes and suggesting these microbiomes can confer benefits to tree saplings. Notably, however, the best model retained the primary axis of functional gene variation of the microbiomes rather than their island type (constructed versus natural). Therefore, although constructed and natural island microbiomes differed in their overall functional composition, it was variation in microbial functional repertoires, rather than island type itself, that best predicted subsequent sapling growth. After filtering for functional genes most relevant to the primary axis of variation (see Methods: *Data analysis*), we manually explored their function and found many of these genes were important for general cellular and carbon metabolism, but interestingly, the genes involved in nitrogen cycling (K00284, K01428, and K00265) were all negatively correlated to the primary axis (Table S2). Additionally, K01626 was also predictive of and negatively correlated to the primary axis of variation in functional gene composition, and is ultimately required for synthesis of siderophores that may also be important in nitrogen cycling (Bellenger et al., 2008; Y. Zhang et al., 2025). Together, these results support functional capacity for nutrient cycling as an important factor differentiating functional genetic repertoires across islands in ways that are relevant to tree sapling growth.

## Discussion

Overall, constructed and natural tree island microbial communities demonstrated robust differences in their functional genetic profiles, corresponding to differences in taxonomic community composition. These differences indicate strong and persistent divergence of microbiomes in constructed habitats from their natural counterparts even over multiple decades. In particular, constructed tree island microbiomes were enriched for functional genes collectively consistent with habitat disturbance, and despite the immediate benefits they conferred during sapling establishment, exhibited reduced functional genetic diversity and redundancy that may simultaneously diminish their long-term health. Because functional genetic diversity and redundancy can help maintain functional stability under variable conditions or environmental regime shifts, constructed islands may lack some capacity for flexibility and resilience that natural tree island microbial communities have, with crucial consequences for long-term restoration and microbial function in such a dynamic and human-impacted ecosystem.

### Taxonomic divergence of microbial communities in reconstructed habitats is reflected in differences in functional repertoires

We found that soil microbial communities in constructed tree islands not only differed from those in natural islands in terms of taxonomic composition (aligning with a previous ITS amplicon analysis of soil fungal communities; Kiesewetter et al., 2025), but also in terms of functional genetic repertoires (i.e., composition when considering both relative abundance and presence/absence metrics; Figures 1 and 2). This is particularly notable because microbial communities are often hyperdiverse, allowing for high degrees of functional redundancy (Afkhami et al., 2026; Louca et al., 2018). Thus, in many systems, habitats maintain similar functional repertoires despite strong taxonomic differentiation (Barnes et al., 2020; Igwe & Afkhami, in revision; Zeng et al., 2023). In fact, a global analysis of publicly available metagenomes found functional gene redundancy despite widespread variation in taxonomic composition (H. Chen et al., 2022), and a study simulating microbial biodiversity loss in metagenomic data found only limited evidence for shifts in functional gene profiles accompanying simulated changes in community taxonomy (H. Chen et al., 2021). These studies suggest community-level functional repertoires are often robust to taxonomic shifts between habitats in natural ecosystems. Therefore, the differences in both taxonomic profiles and functional repertoires between constructed and natural Everglades tree island microbial communities reveal striking and robust divergence of microbial communities in reconstructed habitat, even nearly two decades post-restoration.

### Constructed tree island microbiomes harbor signatures of altered metabolic pathways, disturbance, and lowered capacity for resilience

Constructed tree island microbiomes were enriched in functional pathways related to lipid metabolism, pollutant degradation, and pathogenicity and antibiotic resistance, whereas natural island microbiomes were enriched in functional pathways for general cell maintenance and carbohydrate metabolism. Interestingly, restoration has the potential to alter chemical cycling in microbial communities (Hu et al., 2024; S. Zhang et al., 2024), and this is reflected in the comparative enrichment for lipid metabolism and depletion in carbohydrate metabolism pathways that we identified in constructed island microbiomes. This in turn has implications for carbon cycling and storage, because lipids are relatively recalcitrant in soil (Lorenz et al., 2007) and serve as important storage molecules that can help stabilize soil organic carbon (Rempfert et al., 2026). Therefore, the enrichment of pathways involved in lipid metabolism and degradation in constructed tree island microbial communities indicates potential impacts on broader carbon loss from soils and, ultimately, climate change (Schmitz & Sylvén, 2023). Long-term recovery may eventually allow for microbial functional capacity in restored habitat to approach that in natural habitats, even if community taxonomic compositional differences persist (S. Zhang et al., 2024), but a microbially-informed approach to restoration may be valuable in supporting more robust soil organic carbon storage (e.g., Oberle et al., 2022). Taken together, we found microbiomes in reconstructed habitats exhibit shifts in carbon metabolic potential in ways that could impact broader carbon sequestration and cycling, which could be expanded on with further investigation of resource and substrate utilization.

Our GSEA also provides evidence of anthropogenic disturbance in constructed tree islands. For example, constructed tree island microbiomes were enriched in many pathways for pollutant (e.g., fluorobenzate, styrene) degradation. This is notable because microbes play an important role in bioremediation and degradation of pollution (Fuentes et al., 2014), and the presence of human pollutants can select for microbes with the ability to degrade them (Campeão et al., 2017; Chakraborty & Das, 2016; van der Meer, 2006). Thus, the enrichment of pollutant degradation pathways indicates restored habitats may maintain elevated levels of pollutants relative to natural habitats that in turn exert a selective pressure on microbial communities. Interestingly, one study found that oil contamination led to not only an increase in oil-degrading microbial taxa, but also an increase in lipid and fatty acid metabolism alongside a concomitant decrease in carbohydrate metabolism (Campeão et al., 2017). This aligns with the enrichment in pathways for lipid metabolism relative to carbohydrate metabolism we identified in constructed tree islands and indicates this pattern may result from the availability of alternative anthropogenically-derived carbon sources in the environment.

Additionally, our GSEA identified enrichment of pathways related to pathogenicity and antibiotic resistance (e.g., mycolic acid and lipoarabinomannan biosynthesis) in constructed island microbiomes. Interestingly, this is consistent with predictions from our previous ITS amplicon-based findings (Kiesewetter et al., 2025; see Appendix: Section S2 *Relative enrichment in pathogenicity and antibiotic resistance pathways in constructed tree island microbiomes is consistent with previous ITS-based findings* for details). The enrichment of pathogenicity and antibiotic resistance genes we detected in constructed islands often result from antibiotic contaminants (which may be residual from human activities such as agriculture) exerting a selective pressure on microbial communities (Naga et al., 2025). Other disturbances, including herbicide application, may also select for antibiotic resistance genes in microbiomes (Knecht et al., 2026; Liao et al., 2021). In alignment with this, the constructed islands we sampled from were previously an agricultural site and were subject to repeated herbicide application during early restoration (Aich et al., 2011; E. Cline, personal communication, July 2026). This demonstrates that historical habitat degradation and the subsequent restoration process itself may introduce disturbances that leave legacies, such as enrichment in pathogenicity and antibiotic resistance pathways, and in severe cases even induce a degree of microbial community dysbiosis, with negative consequences for plant fitness (Ketehouli, Goss, et al., 2024; Ketehouli, Pasche, et al., 2024). Therefore, active management of microbial communities may be helpful in overcoming barriers related to historical context, promoting microbiome health, and improving restoration outcomes. Notably, both enrichment for pathogenicity and antibiotic resistance pathways and the enrichment for pollution degradation pathways described above may reflect anthropogenic disturbance, because pollution and microbial pathogenicity may be linked; antibiotic-resistant, opportunistic pathogens are often found in human-disturbed environments due to their ability to adapt to various ecosystems, including those that are human-impacted (Djouadi et al., 2017). In fact, pollutant-contaminated water could serve as a pathogen reservoir through pollutants simultaneously selecting for pollutant degradation genes and enriching for pathogenic taxa (X. Lin et al., 2023). Furthermore, co-contamination of antibiotics and other pollutants often occurs in ecosystems, and pollutants such as heavy metals can even induce antibiotic resistance in microbes (S. Chen et al., 2015), reinforcing this link between environmental disturbance and microbiome antibiotic resistance and pathogenicity. Taken together, our assessment of functional pathways in constructed tree island microbiomes reveals shifts in their functional repertoires that reflect differential environmental disturbance regimes, with possible downstream implications for human health (Naga et al., 2025).

Our greenhouse experiment demonstrated microbiomes with more negative values along the primary axis of variation in functional gene composition conferred greater benefits to tree saplings (Figure 5). Notably, this axis of variation was strongly negatively correlated to as well as predicted by multiple functional genes involved in nitrogen metabolism and assimilation (Table S2). This suggests microbiomes with more negative values along this axis have greater functional genetic capacity for nitrogen cycling, and plants in turn depend on microbial nitrogen cycling for assimilation of this often limiting nutrient and therefore growth (Moreau et al., 2019). In applied settings, microbiome-based management has shown promise and the inoculation of whole microbiomes from desired habitats has contributed to the successful restoration of plant communities (Crawford et al., 2020; Wubs et al., 2016). Building off this, targeted inoculation of communities with the capacity for desired functions such as nitrogen cycling could further improve restoration outcomes. Additionally, because constructed tree island microbiomes in our study generally had more negative values along the primary axis of variation in functional gene composition, they may have a higher abundance of these important nitrogen cycling genes and thus be more effective in promoting sapling growth, demonstrating how natural states do not always represent the most immediate pathway to achieving restoration goals in complex, human-altered ecosystems (Hart et al., 2020).

However, restored ecosystems sometimes exhibit reduced biodiversity and altered abiotic conditions that could weaken long-term success and resilience (Loisel & Gallego-Sala, 2022). Our results show that constructed tree island microbiomes simultaneously exhibited several characteristics that may indicate disturbance and reduced resilience relative to natural island microbiomes. For example, constructed island microbiomes had lower diversity of functional genes than natural island microbiomes, indicating reduced functional capacity, despite no detectable differences in taxonomic diversity. Because keystone taxa allow for unique functions either directly or indirectly via interactions with other microbes (Afkhami et al., 2026), reduced diversity in functional abilities in constructed island microbiomes could potentially reflect missing putative keystone taxa previously identified using ITS amplicon and co-occurrence network-based approaches (Kiesewetter et al., 2025). Importantly, keystone taxa identified using this approach have been shown to drive microbial community reassembly following disturbance (Rawstern et al., 2025). Thus, their reduced relative presence in reconstructed habitat could have long-term consequences for microbiomes such as the reduced functional diversity we identified here. Moreover, because these putative keystone microbial taxa often have narrower abiotic niches (Hernandez et al., 2023), their reduced abundance provides another line of support implicating increased disturbance as a structuring feature in these reconstructed habitats, which is further consistent with the differential enrichment for functional pathways indicative of disturbance identified through our GSEA. We also found reduced functional pathway redundancy in constructed tree island microbiomes, suggesting that microbiome functions may be vulnerable to stress. Essentially, if stress removes taxa with functional abilities that are important, but have low redundancy, lower community resilience is expected (Afkhami et al., 2026). Taken together, the greater functional gene diversity (and thus functional capacity) and elevated functional pathway redundancy (i.e., more functions are able to be carried out by an increased number of microbial taxa) in natural tree island soil microbiomes reflects a comparatively greater potential for functional flexibility and context-dependency, which is crucial for matching environmental conditions in an increasingly stressful and unpredictable world (Hernandez et al., 2025). This highlights the possibility of trade-offs in immediate function and resilience, wherein constructed tree island microbial communities have high capacity for functions (such as those related to nitrogen cycling) that enable them to support increased tree sapling growth, but are also less functionally resilient and therefore more vulnerable to disturbance. Long-term monitoring of constructed and natural tree island microbial communities will be a crucial next step in evaluating resilience and vulnerabilities of these microbiomes to environmental stress, as well as whether constructed island microbiomes will with time converge towards natural islands in terms of their capacity for community resilience. Furthermore, our research emphasizes the importance of future work coupling long-term microbiome monitoring with experimental manipulation of microbial inoculation during tree island construction to develop a holistic understanding of microbial roles in and responses to reconstructive restoration and ecosystem recovery.

Long-term passive restoration may allow for biodiversity and abiotic conditions to eventually return to non-impacted states (Fanelli et al., 2023), although active management may also aid in ecosystem recovery (Clements et al., 2010). Active management may be particularly important if the act of reconstructive restoration itself introduces human disturbance. For instance, herbicide application to remove undesired vegetation may select for microbial antibiotic resistance genes (Knecht et al., 2026; Liao et al., 2021). Additionally, environmental contaminants may accumulate at the bottom of water bodies (Zonta et al., 2020), and soils derived from dredged sediments may contain elevated levels of pollutants (Vandecasteele et al., 2004). Thus, habitat features reconstructed using dredged substrate and were subject to herbicide application, including constructed tree islands (Aich et al., 2011; E. Cline, personal communication, July 2026), may contain substances that in turn continue to shape microbial communities and their functional pathways (such as those for pollutant degradation and antibiotic resistance), even nearly two decades post-construction. Ultimately, more time as well as deliberate management of the soil microbiome may help bring functional profiles of these hidden communities in reconstructed landscapes in line with those in natural habitats.

### Long-term implications of reconstructive restoration for microbiome function

Even nearly two decades post-restoration, microbial communities in constructed tree islands continued to show differences in both taxonomic and functional gene composition from their natural island counterparts, as well as lowered functional genetic diversity and redundancy, with implications for higher-order effects on tree sapling growth. These taxonomic and functional profile differences could be the outcome of dispersal limitation or differences in environmental drivers and filtering (W. Chen et al., 2020). In our system, the similar climatic and hydrological conditions between constructed and natural tree islands could implicate dispersal limitations maintaining divergence between these microbial communities, particularly given the islands are separated by a water matrix and the slow flow of water in the Everglades (Kushlan, 1991). However, our findings here regarding microbiome functional repertoires support that environmental filtering could also play a key role in driving differences in functional profiles between constructed and natural tree island microbiomes. A previous study of this system found constructed tree island communities are depleted in taxa central in networks (Kiesewetter et al., 2025), which usually have narrow abiotic niches (Hernandez et al., 2023). The selective removal of taxa with narrow niches suggests environmental filtering rather than dispersal limitation may play a strong role in determining community composition (W. Chen et al., 2020). Aligning with this, our evaluation of soil metagenomes found evidence of potential long-term disturbances or environmental factors, such as enrichment of pollutant degradation pathways that may indicate the presence of pollutants (Chakraborty & Das, 2016; van der Meer, 2006), in reconstructed habitats that continue to exert a selective influence on microbial communities. This suggests that there may be an inherent and persistent impact of the act of restoration itself on microbial communities, creating lasting legacies in microbiomes that may need time or more active remediation to address and to bring their functional capacities in line with those of natural communities. Overall, we reveal that there are long-term consequences such as lower diversity and redundancy of functional genes and pathways, respectively, when allowing microbial communities to assemble freely in reconstructed ecosystems. Given the importance and ubiquitousness of soil microbes (Hartmann & Six, 2023; Wang et al., 2024), our work demonstrates the value of a microbial component to ecological management (such as inoculation of microbiomes with desired capacities for functions such as nitrogen cycling, or alleviating environmental pressures found to shift microbiome functional profiles), as well as the potential of microbially-informed management decisions (such as leveraging microbial data as indicators of reconstructed habitat health and quality), for more effective and holistic restoration of ecosystem health.

## Data availability

All experimental data and scrips and full details regarding model selection are available from Zenodo (https://doi.org/10.5281/zenodo.22117662). Demultiplexed sequencing data and associated metadata will be available in the National Center for Biotechnology Information (NCBI) Sequence Read Archive under accession number upon acceptance of this manuscript.

## Author Contributions

VWL performed sample preparation for sequencing, bioinformatics processing, statistical analysis, and manuscript writing. ALR contributed to sample preparation for sequencing and bioinformatics processing. KNK collected field samples and set up and managed the experiment, as well as contributed to study conceptualization and design and statistical analysis. EC managed and provided access to constructed tree islands and contributed to study conceptualization and design. AHR contributed to bioinformatics processing, statistical analysis, and setting up the experiment. CC contributed to study conceptualization and design, field sample collection, and providing access to natural tree islands. FHS contributed to study conceptualization and design, field sample collection, and providing access to tree islands. MEA contributed to setting up the experiment and led study conceptualization and design, funding acquisition, establishing a team of collaborators, and student supervision. All authors contributed to manuscript editing and revisions.

## Acknowledgements

We thank Fabiola Santamaria for her help in field sample collection and providing access to natural tree islands, Damian Hernandez and Brianna Almeida for their help setting up the greenhouse experiment, Gina R. Ortiz for procuring/repotting tree saplings and providing care during acclimation prior to the greenhouse experiment, and J.E. Cary, O.D. Morales Casanova, K. Kirejevas, L.J. Carbajal, and G.B. Pohlmann for their help with plant care, data collection, and data input. We also thank Jordan Busch for his help in maintaining the saplings prior to the experiment. Additionally, we thank Caitlin Broderick for her very helpful bioinformatics advice. This study was primarily funded by the South Florida Water Management District to MEA with additional support from NSF DEB-1922521 and NSF DEB-2030060 to MEA, the NSF Graduate Research Fellowship Program and University of Miami Dissertation Year Fellowship to KNK, and the USDA NIFA Predoctoral Fellowship to AHR.

