## Appendix for "Hidden drivers of restoration: Persistent divergence in soil microbiome functional capacity post-habitat reconstruction"

### Appendix 1: Supporting Information Contents

Section S1: Supplementary Methods ………………………………………………… Pages 1 – 2

Section S2: Supplementary Discussion …………………………………………………… Page 2

Supplementary Figures (S1) ….…………………………………………………………… Page 3

Supplementary Tables (S1–S16) .……………………………………………………. Pages 3 – 12

Supplementary References ………………………………………………………… Pages 13 – 16

#### Section S1: Supplementary Methods

##### Manipulation of hydrological treatments in the plant–microbiome experiment

Hydrological treatments applied in our plant–microbiome greenhouse experiment simulated different management practices being considered for the Everglades. The “unconstrained” hydrological regime allows natural water accumulation from precipitation, often leading to inundation of tree islands during the wet season. The “constrained” hydrological regime limits water depth through diversion of water into canals (Ask IFAS - Powered by EDIS, n.d.), thereby preventing inundation of tree islands during the wet season. Using data available from LILA, we simulated both these management practices. To do so, we manipulated water stage to match projected unconstrained and constrained stages during the wet part of the hydrograph relative to average tree island soil surface height (which we treated the soil surface of microcosms as proxies for) and the height of water at the lowest water level during the year (which we treated the bottoms of microcosm vessels as proxies for). To calculate the exact water heights for our experimental treatments, we then determined the proportion of that height difference that would have been flooded for each management practice based on the projected water stage for an island in the field in December, a good model for wet period hydrology (see Kiesewetter et al. 2025). These treatments were then applied by watering all microcosms the same amount from above (to simulate equal precipitation), and then allowing excess water to drain through holes drilled in the sides of outer vessels for the constrained treatment (to simulate diverting water).

##### Constructed and natural tree island microbiomes show robust differences in functional genetic redundancy

After evaluating functional redundancy for each KEGG functional pathway within each sample using Kaiju taxonomic assignments to the genus level, we employed a paired design when testing for differences in functional redundancy between constructed and natural islands. Specifically, differences in functional redundancy were evaluated using a paired t-test comparing the difference in mean functional redundancy across constructed islands compared to the mean functional redundancy across natural islands for each given KEGG functional pathway (see Methods section). However, we were concerned some functional pathways may have low representation across constructed and/or natural samples, making their mean functional redundancy more susceptible to influence from a small number of samples relative to other, more abundant functional pathways.

To explore and account for this possibility, we employed several alternative approaches. First, we replicated the analysis described in the main text, but excluded any functional pathways that occurred in <25% of either constructed or natural samples. In doing so, we removed pathways with functional redundancy values representing an average of very few samples that may be strongly influenced by individual sample values. This led to a filtering of five pathways, with 431 retained in our analysis. Following this additional filtering, we found that natural tree island microbial communities had relatively higher functional redundancy (paired t-test: t = 8.59, df = 430, p < 0.0001; mean ± S.E.M. difference in functional redundancy for a given functional pathway: 0.11 ± 0.01; n = 431 pathways). We further used a linear mixed-effects model with functional pathway as a random effect to test how functional redundancy depended on island type (constructed/natural) using *lmer* (*lme4*; Bates et al. 2026), which identified natural island microbiomes as having higher functional pathway redundancy (χ^2^_1_ = 112.68, df = 1, p < 0.0001). When using a linear model with functional pathway as a fixed effect instead, we again identified natural island microbiomes as having higher functional pathway redundancy (F_1,6694_ = 112.03, p < 0.0001). Overall, the finding that natural island microbiomes had greater functional redundancy than constructed island microbiomes was robust to analytical decisions.

#### Section S2: Supplementary Discussion

##### Relative enrichment in pathogenicity and antibiotic resistance pathways in constructed tree island microbiomes is consistent with previous ITS-based findings

Our GSEA identified significant enrichment of many pathways related to pathogenicity and antibiotic resistance in constructed island microbiomes (Figure 4). This aligns with predictions from our previous ITS amplicon-based findings, where the majority of fungal taxa predictive of constructed habitats were identified as potential pathogens (Kiesewetter et al. 2025). Notably, however, by directly annotating functional genes, the data presented here more conclusively identify pathogenic capacity within a microbiome than previous approaches relying on matching taxonomic IDs to hypothesized functional guilds in databases, which may not be able to match the majority of taxa and cannot account for context-dependency of a taxa’s functional roles or variation between microbial strains (Kiesewetter et al. 2025). Nonetheless, the consistency of functional characterization between approaches supports the utility of amplicon analysis for rough insight into the functional attributes of a community.

#### Supplementary Figures

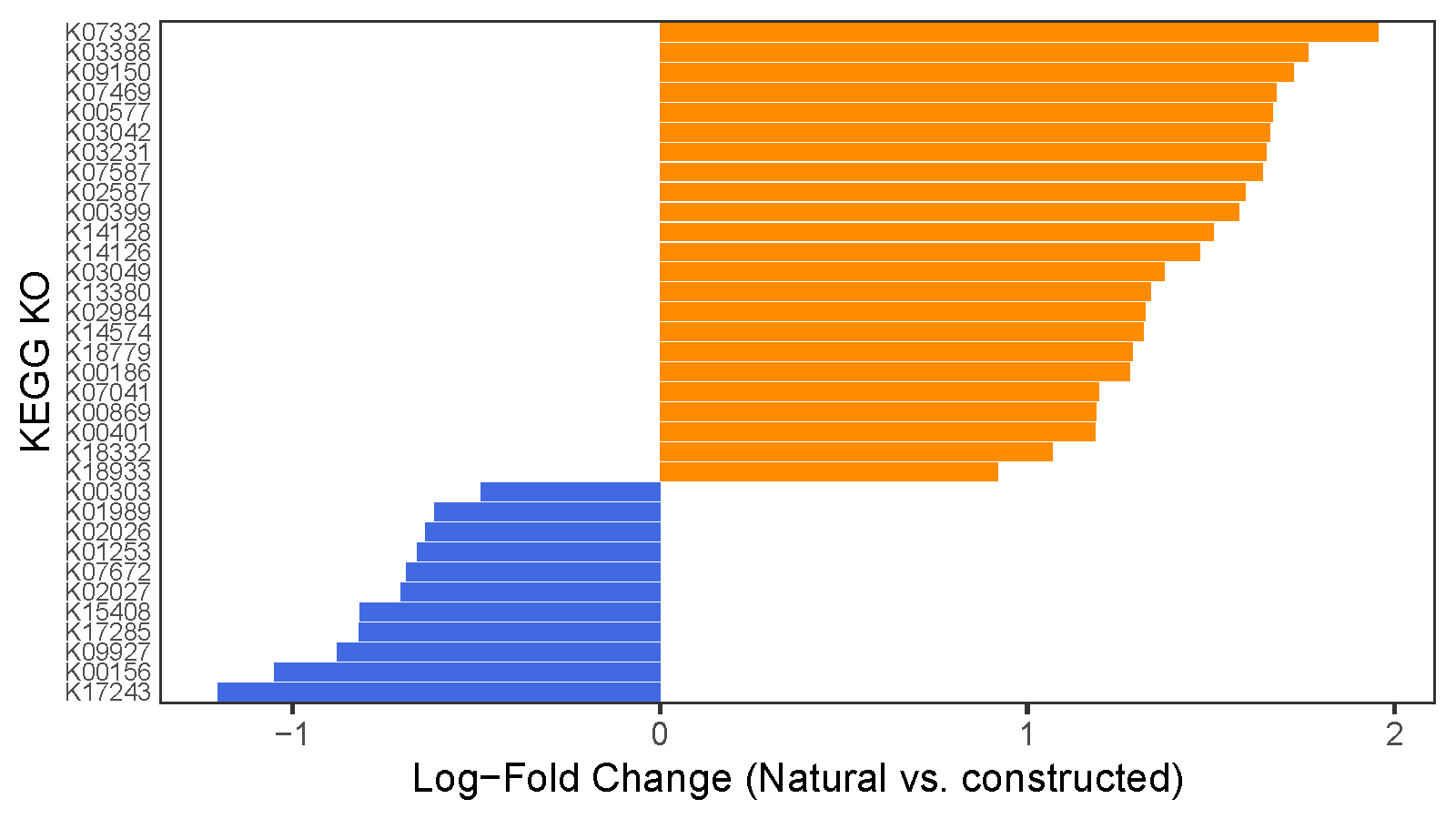

**Figure S1.** KEGG KOs that exhibited significantly differential abundance between constructed and natural tree island soil microbiomes, identified by ANCOM-BC2. KEGG KOs with positive log-fold change values were relatively more abundant in natural island microbiomes, whereas those with negative log-fold change values were more abundant in constructed island microbiomes.

#### Supplementary Tables

**Table S1.** GPS locations of the 14 natural islands within Water Conservation Area 3A where soils were collected.

| **Island ID** | **Location** | **Latitude** | **Longitutde** |
| --- | --- | --- | --- |
| 1 | Head | N 25^o^47’45.3” | W 80^o^41’22.2” |
| 1 | Tail | N 25^o^46’52.7” | W 80^o^41’20.2” |
| 2 | Head | N 25^o^47’46.2” | W 80^o^41’21.5” |
| 2 | Tail | N 25^o^47’45.3” | W 80^o^41’22.2” |
| *3* | *Head* | *N 25^o^49’13.4”* | *W 80^o^41’52.1”* |
| 4 | Head | N 25^o^52’52.9” | W 80^o^40’59.1” |
| 4 | Tail | N 25^o^52’47.3” | W 80^o^40’58.9” |
| 5 | Head | N 25^o^54’49.9” | W 80^o^39’19.7” |
| 5 | Tail | N 25^o^54’46.9” | W 80^o^39’21.3” |
| 6 | Head | N 25^o^53’48.0” | W 80^o^41’02.6” |
| 6 | Tail | N 25^o^53’45.6” | W 80^o^41’02.8” |
| 7 | Head | N 25^o^52’17.8” | W 80^o^41’56.4” |
| 7 | Tail | N 25^o^52’16.3” | W 80^o^41’57.2” |
| 8 | Head | N 25^o^51’24.3” | W 80^o^41’51.3” |
| 8 | Tail | N 25^o^51’22.2” | W 80^o^41’53.6” |
| *10* | *Head* | *N 25^o^46’18.2”* | *W 80^o^44’09.2”* |
| *10* | *Tail* | *N 25^o^46’18.2”* | *W 80^o^44’09.2”* |
| *11* | *Head* | *N 25^o^46’52.2”* | *W 80^o^44’30.0”* |
| *11* | *Tail* | *N 25^o^46’49.8”* | *W 80^o^44’29.8”* |
| *12* | *Head* | *N 25^o^47’59.8”* | *W 80^o^44’38.8”* |
| *12* | *Tail* | *N 25^o^47’57.8”* | *W 80^o^44’39.2”* |
| *13* | *Head* | *N 25^o^48’15.3”* | *W 80^o^45’19.2”* |
| *13* | *Tail* | *N 25^o^48’11.0”* | *W 80^o^45’21.4”* |
| *14* | *Head* | *N 25^o^49’06.0”* | *W 80^o^45’13.7”* |
| *14* | *Tail* | *N 25^o^49’04.3”* | *W 80^o^45’15.1”* |
| *15* | *Head* | *N 25^o^51’23.6”* | *W 80^o^46’10.6”* |
| *15* | *Tail* | *N 25^o^51’22.8”* | *W 80^o^46’11.1”* |
| NOTE: Islands in italics are those used both for microbiome sequencing and the greenhouse experiments. | | | |

**Table S2.** KEGG KOs identified using Boruta feature selection as predictive of the primary axis of variation in functional gene variation by Aitchison distance matrices, following filtering to retain only those that were highly significant, strongly correlated to the primary axis, and well-explained overall by our PCoA.

| **KEGG KO** | **Direction of correlation** | **r** | **Name** | **Function** | **Citation** |
| --- | --- | --- | --- | --- | --- |
| K02843 | Positive | >0.99 | lipopolysaccharide heptosyltransferase II [EC:2.4.99.24] | Enzyme for lipopolysaccharide synthesis, which is crucial for bacterial survival | (Gronow et al. 2000) |
| K07798 | Positive | 0.92 | membrane fusion protein, copper/silver efflux system | Periplasmic protein involved in copper and silver efflux, important for metal tolerance | (Bagai et al. 2007) |
| K19113 | Positive | 0.87 | acetyltransferase [EC:2.3.1.-] | Enzyme involved in enabling pathogens to infect plants | (Wencewicz and Walsh 2012) |
| K00334 | Positive | 0.86 | NADH-quinone oxidoreductase subunit E [EC:7.1.1.2] | Subunit of mitochondrial NADH-quinone oxidoreductase, needed for energy transduction | (Weidner et al. 1993) |
| K16887 | Positive | 0.85 | quinone-modifying oxidoreductase, subunit QmoC | Subunit of enzyme putatively involved in sulfur metabolism | (Falkenby et al. 2011) |
| K09516 | Positive | 0.84 | all-trans-retinol 13,14-reductase [EC:1.3.99.23] | Enzyme for retinol metabolism; produces metabolite with unknown function | (Moise et al. 2004) |
| K14534 | Positive | 0.84 | 4-hydroxybutyryl-CoA dehydratase / vinylacetyl-CoA-Delta-isomerase [EC:4.2.1.120 5.3.3.3] | Enzyme for metabolism of aminobutyrate as a carbon source | (Scherf and Buckel 1993) |
| K19664 | Positive | 0.83 | CDP-2,3-bis-(O-geranylgeranyl)-sn-glycerol synthase [EC:2.7.7.67] | Enzyme for archaeal lipid synthesis | (Jain et al. 2014) |
| K09835 | Positive | 0.82 | prolycopene isomerase [EC:5.2.1.13] | Enzyme for biosynthesis of carotenoids, which are essential photoprotective and antioxidant pigments | (Park et al. 2002) |
| K17992 | Positive | 0.82 | NADP-reducing hydrogenase subunit HndB [EC:1.12.1.3] | Subunit of NADP-reducing hydrogenase, important for metabolism of sulfate-reducing bacteria | (de Luca et al. 1998) |
| K02626 | Positive | 0.82 | arginine decarboxylase [EC:4.1.1.19] | Enzyme for arginine metabolism and polyamine biosynthesis; involved in bacterial acid resistance | (Giles and Graham 2007; Graham et al. 2002) |
| K02548 | Positive | 0.82 | 1,4-dihydroxy-2-naphthoate polyprenyltransferase [EC:2.5.1.74] | Enzyme for biosynthesis of menaquinone (vitamin K2), which is crucial for bacterial growth and survival | (Suvarna et al. 1998) |
| K13819 | Positive | 0.81 | NifU-like protein | Largely unclassified |  |
| K03750 | Positive | 0.80 | molybdopterin molybdotransferase [EC:2.10.1.1] | Enzyme for biosynthesis of molybdenum cofactors, required for survival of most bacteria | (Nichols and Rajagopalan 2005) |
| K04770 | Positive | 0.80 | Lon-like ATP-dependent protease [EC:3.4.21.-] | Enzyme involved in protein metabolism | (Papanastasiou et al. 2013) |
| K14654 | Positive | 0.74 | 2,5-diamino-6-(ribosylamino)-4(3H)-pyrimidinone 5'-phosphate reductase [EC:1.1.1.302] | Enzyme involved in biosynthesis of riboflavin (vitamin B2), which is essential for cell reactions | (Averianova et al. 2020; Chatwell et al. 2006) |
| K11261 | Positive | 0.73 | formylmethanofuran dehydrogenase subunit E [EC:1.2.7.12] | Subunit of enzyme for methanogenesis (and thus decomposition) | (Vorholt et al. 1996) |
| K02978 | Positive | 0.71 | small subunit ribosomal protein S27e | Ribosomal subunit protein | (Chan et al. 1993) |
| K00284 | Negative | –0.79 | glutamate synthase (ferredoxin) [EC:1.4.7.1] | Enzyme important for nitrogen (ammonium) assimilation | (Kameya et al. 2007; Vanoni and Curti 1999) |
| K02274 | Negative | –0.84 | cytochrome c oxidase subunit I [EC:7.1.1.9] | Enzyme required for heterotrophic growth | (Schmetterer et al. 2001) |
| K00164 | Negative | –0.84 | 2-oxoglutarate dehydrogenase E1 component [EC:1.2.4.2] | Component of enzyme important for cell metabolism | (Repetto and Tzagoloff 1989) |
| K01638 | Negative | –0.84 | malate synthase [EC:2.3.3.9] | Enzyme for converting lipids to sugars, important for carbon metabolism | (Molina et al. 1994) |
| K00128 | Negative | –0.85 | aldehyde dehydrogenase (NAD+) [EC:1.2.1.3] | Enzyme important for cell metabolism | (Racker 1949) |
| K01915 | Negative | –0.88 | glutamine synthetase [EC:6.3.1.2] | Enzyme for glutamine biosynthesis; important for cell metabolism | (Newsholme et al. 2003) |
| K01681 | Negative | –0.89 | aconitate hydratase [EC:4.2.1.3] | Enzyme for cell respiratory metabolism | (Kesawat et al. 2022) |
| K01626 | Negative | –0.91 | 3-deoxy-7-phosphoheptulonate synthase [EC:2.5.1.54] | Enzyme needed for the shikimate pathway, which is responsible for biosynthesis of chorismite; chorismite is the precursor of many aromatic compounds such as many amino acids and salicylate (required for biosynthesis of siderophore for iron acquisition) | (Webby et al. 2005) |
| K01428 | Negative | –0.91 | urease subunit alpha [EC:3.5.1.5] | Subunit of urease, an enzyme for urea degradation that is important for nitrogen cycling and acid resistance | (Cruz-Ramos et al. 1997; Zhou et al. 2019) |
| K00549 | Negative | –0.91 | 5-methyltetrahydropteroyltriglutamate--homocysteine methyltransferase [EC:2.1.1.14] | Enzyme for amino acid biosynthesis | (Gonzalez et al. 1992) |
| K00163 | Negative | –0.92 | pyruvate dehydrogenase E1 component [EC:1.2.4.1] | Component of enzyme important for cell metabolism | (Tian et al. 2005) |
| K00265 | Negative | –0.94 | glutamate synthase (NADPH) large chain [EC:1.4.1.13] | Major subunit of glutamate synthase, an enzyme important for nitrogen (ammonium) assimilation | (Sonawane and Röhm 2004; Vanoni and Curti 1999) |

**Table S3. Taxonomic composition using robust Aitchison distance metrics as a function of island source and location.** PERMANOVA statistical results for soil microbiome taxonomic composition at the genus level using robust Aitchison distance metrics as a function of island type (constructed/natural) and location (head/tail), using 999 permutations.

|  | F | DF | P |
| --- | --- | --- | --- |
| Source | 13.29 | 1, 39 | 0.001 |
| Location | 2.06 | 1, 39 | 0.12 |

**Table S4. Taxonomic composition using Jaccard distance metrics as a function of island source and location.** PERMANOVA statistical results for soil microbiome taxonomic composition at the genus level using Jaccard distance metrics as a function of island type (constructed/natural) and location (head/tail), using 999 permutations.

|  | F | DF | P |
| --- | --- | --- | --- |
| Source | 1.99 | 1, 39 | 0.002 |
| Location | 1.03 | 1, 39 | 0.27 |

**Table S5. Taxonomic Shannon diversity as a function of island source and location.** ANOVA statistical results for soil microbiome taxonomic diversity at the genus level as a function of island type (constructed/natural) and location (head/tail).

|  | χ^2^ | DF | P |
| --- | --- | --- | --- |
| Source | 0.44 | 1 | 0.51 |
| Location | 1.15 | 1 | 0.28 |

**Table S6. Taxonomic richness as a function of island source and location.** ANOVA statistical results for soil microbiome taxonomic richness at the genus level as a function of island type (constructed/natural) and location (head/tail).

|  | χ^2^ | DF | P |
| --- | --- | --- | --- |
| Source | 0.62 | 1 | 0.43 |
| Location | 1.83 | 1 | 0.18 |

**Table S7. Functional genetic composition using robust Aitchison distance metrics as a function of island source and location.** PERMANOVA statistical results for soil microbiome functional genetic composition using KEGG KOs and robust Aitchison distance metrics as a function of island type (constructed/natural) and location (head/tail), using 999 permutations.

|  | F | DF | P |
| --- | --- | --- | --- |
| Source | 14.03 | 1, 39 | 0.001 |
| Location | 0.49 | 1, 39 | 0.60 |

**Table S8. Functional genetic composition using Jaccard distance metrics as a function of island source and location.** PERMANOVA statistical results for soil microbiome functional genetic composition using KEGG KOs and Jaccard distance metrics as a function of island type (constructed/natural) and location (head/tail), using 999 permutations.

|  | F | DF | P |
| --- | --- | --- | --- |
| Source | 3.29 | 1, 39 | 0.001 |
| Location | 0.98 | 1, 39 | 0.41 |

**Table S9. Functional genetic Shannon diversity as a function of island source and location.** ANOVA statistical results for soil microbiome functional genetic diversity using KEGG KOs as a function of island type (constructed/natural) and location (head/tail).

|  | χ^2^ | DF | P |
| --- | --- | --- | --- |
| Source | 5.65 | 1 | 0.02 |
| Location | 0.62 | 1 | 0.43 |

**Table S10. Functional genetic richness as a function of island source and location.** ANOVA statistical results for soil microbiome functional genetic richness using KEGG KOs as a function of island type (constructed/natural) and location (head/tail).

|  | χ^2^ | DF | P |
| --- | --- | --- | --- |
| Source | 0.01 | 1 | 0.91 |
| Location | 0.12 | 1 | 0.73 |

**Table S11. Functional genetic redundancy as a function of island source.** Paired t-test for differences in redundancy of KEGG functional pathways between constructed and natural tree island soil microbiomes.

|  | t | DF | P |
| --- | --- | --- | --- |
| Source | 8.63 | 1, 432 | <0.001 |

**Table S12.** Significantly enriched KEGG functional pathways based on gene set enrichment analysis.

| KEGG pathway | Description | Enrichment score | p_adj_ |
| --- | --- | --- | --- |
| ko00680 | Methane metabolism | 0.704577 | 3.72E-09 |
| ko05171 | Coronavirus disease - COVID-19 | 0.878072 | 3.72E-09 |
| ko03010 | Ribosome | 0.701941 | 3.72E-09 |
| ko01200 | Carbon metabolism | 0.531656 | 3.72E-09 |
| ko02010 | ABC transporters | -0.41855 | 3.72E-09 |
| ko03030 | DNA replication | 0.801289 | 1.15E-08 |
| ko03020 | RNA polymerase | 0.850606 | 1.63E-07 |
| ko01220 | Degradation of aromatic compounds | -0.45999 | 2.23E-06 |
| ko00970 | Aminoacyl-tRNA biosynthesis | 0.706647 | 1.59E-05 |
| ko00720 | Other carbon fixation pathways | 0.576538 | 2.56E-05 |
| ko03008 | Ribosome biogenesis in eukaryotes | 0.775407 | 6.72E-05 |
| ko00627 | Aminobenzoate degradation | -0.54699 | 0.000143 |
| ko00860 | Porphyrin metabolism | 0.541282 | 0.000197 |
| ko00074 | Mycolic acid biosynthesis | -0.80616 | 0.000262 |
| ko00572 | Arabinogalactan biosynthesis | -0.83466 | 0.000328 |
| ko00360 | Phenylalanine metabolism | -0.51068 | 0.001094 |
| ko00260 | Glycine, serine and threonine metabolism | -0.44668 | 0.001094 |
| ko00622 | Xylene degradation | -0.65809 | 0.001274 |
| ko00361 | Chlorocyclohexane and chlorobenzene degradation | -0.6902 | 0.001274 |
| ko00380 | Tryptophan metabolism | -0.55292 | 0.001274 |
| ko00350 | Tyrosine metabolism | -0.51143 | 0.001274 |
| ko00053 | Ascorbate and aldarate metabolism | -0.58216 | 0.001716 |
| ko00364 | Fluorobenzoate degradation | -0.7836 | 0.001716 |
| ko04141 | Protein processing in endoplasmic reticulum | 0.810444 | 0.001716 |
| ko00571 | Lipoarabinomannan (LAM) biosynthesis | -0.74514 | 0.002875 |
| ko01240 | Biosynthesis of cofactors | 0.375715 | 0.004279 |
| ko00984 | Steroid degradation | -0.6477 | 0.004666 |
| ko01057 | Biosynthesis of type II polyketide products | -0.72511 | 0.006111 |
| ko00623 | Toluene degradation | -0.60173 | 0.006305 |
| ko00071 | Fatty acid degradation | -0.52653 | 0.00781 |
| ko00561 | Glycerolipid metabolism | -0.51205 | 0.01095 |
| ko01210 | 2-Oxocarboxylic acid metabolism | 0.4724 | 0.013142 |
| ko00633 | Nitrotoluene degradation | 0.713115 | 0.022675 |
| ko00040 | Pentose and glucuronate interconversions | -0.43027 | 0.022792 |
| ko00643 | Styrene degradation | -0.62623 | 0.034717 |
| ko00982 | Drug metabolism | -0.69515 | 0.041986 |
| ko00130 | Ubiquinone and other terpenoid-quinone biosynthesis | -0.44017 | 0.048138 |

**Table S13. Trunk diameter as a function of hydrological treatment and the interaction between inoculation treatment and the primary axis of variation in functional gene composition.** ANOVA statistical results for standardized sapling trunk diameter as a function of hydrological treatment (constrained/unconstrained) and the interaction between inoculation treatment (live/sterile) and the primary axis of variation in microbiome functional gene (KEGG KO) composition (using Aitchison distance matrices). Please see Appendix: Table S15 for model comparison details.

|  | χ^2^ | DF | P |
| --- | --- | --- | --- |
| Hydrological treatment | 17.63 | 1 | <0.0001 |
| Inoculation treatment | 0.62 | 1 | 0.43 |
| Primary axis | 0.54 | 1 | 0.46 |
| Inoculation treatment * primary axis | 7.90 | 1 | 0.005 |

**Table S14. Candidate models for leaf number.** Due to the large number of model comparisons, only the ten models with the lowest AIC_c_ scores are listed. Please see Zenodo (<https://doi.org/10.5281/zenodo.22117662>) for a complete table of candidate models and comparisons.

| Candidate model | df | logLik | AICc | delta | weight |
| --- | --- | --- | --- | --- | --- |
| Leaf number ~ 1 | 3 | –418.958 | 843.9976 | 0 | 0.070905 |
| Leaf number ~ PCo1 | 4 | –418.363 | 844.863 | 0.86546 | 0.045999 |
| Leaf number ~ inoculum status | 4 | –418.544 | 845.2242 | 1.226676 | 0.038398 |
| Leaf number ~ PCo1 + island type | 5 | –417.548 | 845.3011 | 1.303526 | 0.03695 |
| Leaf number ~ island type | 4 | –418.604 | 845.3443 | 1.346779 | 0.03616 |
| Leaf number ~ functional genetic diversity | 4 | –418.67 | 845.4764 | 1.478865 | 0.033849 |
| Leaf number ~ hydrological treatment | 4 | –418.916 | 845.9694 | 1.971815 | 0.026455 |
| Leaf number ~ functional genetic redundancy | 4 | –418.958 | 846.0523 | 2.054775 | 0.02538 |
| Leaf number ~ PCo1 + inoculum status | 5 | –417.948 | 846.1007 | 2.103116 | 0.024774 |
| Leaf number ~ PCo1 + inoculum status * island type | 7 | –415.934 | 846.2551 | 2.257508 | 0.022933 |

**Table S15. Candidate models for trunk diameter.** Due to the large number of model comparisons, only the ten models with the lowest AIC_c_ scores are listed. Please see Zenodo (<https://doi.org/10.5281/zenodo.22117662>) for a complete table of candidate models and comparisons.

| Candidate model | df | logLik | AICc | delta | weight |
| --- | --- | --- | --- | --- | --- |
| Trunk diameter ~ Hydrological treatment + PCo1 * inoculum status | 7 | –400.5594392 | 815.5050854 | 0 | 0.142464743 |
| Trunk diameter ~ inoculum status * (PCo1 + hydrological treatment) | 8 | –400.0436163 | 816.5855026 | 1.080417168 | 0.083003765 |
| Trunk diameter ~ PCo1 * (inoculum status + hydrological treatment) | 8 | –400.299139 | 817.0965478 | 1.591462445 | 0.064287379 |
| Trunk diameter ~ Island type + hydrological treatment + PCo1 * inoculum status | 8 | –400.3535176 | 817.2053052 | 1.700219772 | 0.060884868 |
| Trunk diameter ~ PCo1 * (inoculum status + hydrological treatment) + inoculum status * hydrological treatment | 9 | –399.7835277 | 818.1920555 | 2.686970085 | 0.037173997 |
| Trunk diameter ~ island type + inoculum status * (PCo1 + hydrological treatment) | 9 | –399.8323696 | 818.2897392 | 2.784653835 | 0.035401976 |
| Trunk diameter ~ island type + PCo1 * (inoculum status + hydrological treatment) | 9 | –400.093559 | 818.8121181 | 3.307032693 | 0.027264301 |
| Trunk diameter ~ hydrological treatment + inoculum treatment * functional genetic diversity | 7 | –402.2294472 | 818.8451013 | 3.340015895 | 0.026818357 |
| Trunk diameter ~ PCo1 * inoculum treatment * hydrological treatment | 10 | –399.0805066 | 818.9275638 | 3.422478392 | 0.025735088 |
| Trunk diameter ~ PCo1 * inoculum treatment + island type * hydrological treatment | 9 | –400.2353343 | 819.0956687 | 3.590583264 | 0.023660404 |

**Table S16. Candidate models for stomatal conductance.** Due to the large number of model comparisons, only the ten models with the lowest AIC_c_ scores are listed. Please see Zenodo (<https://doi.org/10.5281/zenodo.22117662>) for a complete table of candidate models and comparisons.

| Candidate model | df | logLik | AICc | delta | weight |
| --- | --- | --- | --- | --- | --- |
| Stomatal conductance ~ 1 | 3 | –382.8069555 | 771.7051658 | 0 | 0.087722918 |
| Stomatal conductance ~ functional genetic redundancy | 4 | –381.9866221 | 772.125916 | 0.420750189 | 0.071080149 |
| Stomatal conductance ~ hydrological treatment | 4 | –382.4544945 | 773.0616608 | 1.35649493 | 0.044519875 |
| Stomatal conductance ~ PCo1 | 4 | –382.5206903 | 773.1940524 | 1.488886545 | 0.04166827 |
| Stomatal conductance ~ hydrological treatment + functional genetic redundancy | 5 | –381.6254066 | 773.4806982 | 1.775532338 | 0.036104482 |
| Stomatal conductance ~ functional genetic diversity | 4 | –382.7200572 | 773.5927861 | 1.887620244 | 0.034136701 |
| Stomatal conductance ~ island type | 4 | –382.7831405 | 773.7189527 | 2.013786905 | 0.032049761 |
| Stomatal conductance ~ inoculum status | 4 | –382.7959748 | 773.7446214 | 2.039455611 | 0.031641052 |
| Stomatal conductance ~ functional genetic redundancy + functional genetic diversity | 5 | –381.9472499 | 774.1243849 | 2.419219076 | 0.026168952 |
| Stomatal conductance ~ inoculum status + functional genetic redundancy | 5 | –381.9778432 | 774.1855715 | 2.480405684 | 0.025380479 |
